# Developmental Changes in Visual Responses to Social Interactions

**DOI:** 10.1101/800532

**Authors:** Jon Walbrin, Ioana Mihai, Julia Landsiedel, Kami Koldewyn

**Author notes:** Corresponding author Jon Walbrin.

## Abstract

Recent evidence demonstrates that a region of the posterior superior temporal sulcus (pSTS) is selective to visually observed social interactions in adults. In contrast, we know comparatively little about neural responses to social interactions in children. Here, we used fMRI to ask whether the pSTS would be ‘tuned’ to social interactions in children at all, and if so, how selectivity might differ from adults. This was investigated not only in the pSTS, but also in socially-tuned regions in neighbouring temporal cortex: extrastriate body area (EBA), face-selective STS (STS-F), fusiform face area (FFA), and temporo-parietal junction (TPJ-M).

Both children and adults showed selectivity to social interaction within right pSTS, while only adults showed selectivity on the left. Adults also showed both more focal and greater selectivity than children (6–12 years) bilaterally. Exploratory sub-group analyses showed that younger children (6–8 years), but not older children (9-12), are less selective than adults on the right, while there was a developmental trend (adults > older > younger) in left pSTS. These results suggest that, over development, the neural response to social interactions is characterized by increasingly more selective, more focal and more bilateral pSTS responses, a process that likely continues into adolescence.

**Highlights:**

- Children show less interaction selectivity in the pSTS than adults
- Adults show bilateral pSTS selectivity, while children are more right-lateralized
- Exploratory findings suggest interaction selectivity in pSTS is more focally tuned in adults

## 1. Introduction

As a deeply social species, humans attend to, and draw social inferences from, a wide array of social cues from both individuals and the interactions that take place between them. Social interactions often carry unique social information (e.g. shared or opposing intentions, cues to relationships, or cues to the relative social status of interactors; Quadflieg & Koldewyn, 2017). The importance of these cues is emphasized by evidence that young children are sensitive to interactive cues and can use aspects of interactive behaviour to make their own social choices. For example, infants (< 18 months) are sensitive to the collaborative intent of an interacting dyad (Fawcett & Gredebäck, 2013; Henderson & Woodward, 2011) and show a preference for interactors who help others (Hamlin et al., 2007). However, processing other types of interactive information, such as recognising the relative social status of two interactors (Brey & Shutts, 2015), inferring the presence of an unseen interaction partner from video footage of an individual (Balas, et al., 2012), and interpreting the communicative intent of interacting point-light figures (Centelles, et al., 2013), undergoes substantial development across childhood.

Recent behavioural work exploring responses to human dyads in adults (e.g. Papeo et al., 2017; Ding et al., 2017; Vestner et al., 2018; Papeo & Abassi, 2019; Papeo & Soto-Faraco, 2019) suggests that dyads that appear to be interacting are processed more similarly to *one* entity than two individuals. In addition, the presence of an interactant can bias the perception of their interaction partner’s emotional expression (Gray et al., 2017), or improve one’s ability to ‘see’ an action presented in noise (Neri et al., 2006) even when the interaction partner is task irrelevant. This evidence, in turn, suggests that the brain may process interactions differently than as a simple combination of multiple bodies, faces, and actions.

Functional magnetic resonance imaging (fMRI) evidence has begun to elucidate the neural basis of perceiving social interactions in adults. Recent work suggests specialised processing for dynamic interactions in the posterior superior temporal sulcus (pSTS) for both point-light figures and moving geometric shapes (i.e. Isik et al., 2017; Walbrin et al., 2018). These findings suggest that the pSTS not only demonstrates selectivity to social interactions (i.e. response to two interacting individuals > two non-interacting individuals), but also differentiates between different types of interactions (i.e. competition and cooperation). Indeed, the pSTS is sensitive to a wide variety of visually presented dynamic (Centelles et al., 2011; Georgescu et al., 2014; Kujala, et al., 2012) as well as static social interaction stimuli (Quadflieg et al., 2015). These results emphasize the important role that this region plays in perceiving and understanding social interactions, but it is also clear that activity in this region does not tell the whole story. Indeed, another line of recent research implicates the complementary functioning of neighbouring extrastriate body area (EBA) in the configural processing of both dynamic (Walbrin & Koldewyn, 2019) and static interacting human dyads (Abassi & Papeo, 2019). Additionally, responses across posterior-temporal regions, including EBA and pSTS, are sensitive to the apparent congruency of an interacting dyad (Quadflieg et al., 2015).

An important extension of this work is to study how these functionally specific responses, especially in the pSTS, emerge and change during development. Previous developmental studies investigating functional pSTS responses to social stimuli have typically focused on faces and bodies, and tend to show weaker responses in children compared to adults (Deen et al., 2017; Ross et al., 2014; Scherf et al., 2007), although this is not always the case (e.g. Golarai et al., 2007). Indeed, the broader STS area is known to undergo substantial structural changes between childhood and adulthood (Bonte et al., 2013; Gogtay et al., 2004; Mills et al., 2014). Therefore, developmental changes in pSTS responses to social interactions seem likely.

To our knowledge, there has been only one developmental study looking at fMRI responses to social interactions. Sapey-Triomphe et al. (2017) compared whole-brain responses to interacting point-light figure dyads with similar, but non-interacting dyads in adults (20+ years), adolescents (13–17 years) and children (8–11 years). Although the pSTS was activated across all three age-groups, no activation differences were shown within the pSTS. Instead, parametric analyses showed a tendency for greater recruitment of fronto-parietal regions, and conversely, lesser recruitment of temporo-occipital regions with increasing age. Importantly, subjects made a social judgement for each trial (i.e. identifying if individuals were ‘acting together or separately’), and as such, the resulting activation reflects not only visual processing but also processing involved in making explicit social inferences.

The present work addresses a similar question to Sapey-Triomphe et al. (2017), but with two crucial methodological differences. First, to allow greater sensitivity to potential differences between groups in the pSTS and other ‘social brain’ regions, we adopt a functionally defined region of interest (ROI) approach for our primary analyses. One advantage of this approach over group-level whole-brain analysis is that it is more robust to inter-subject spatial variability in activation maps (Saxe et al., 2006), an important consideration given previously observed age-related increases in the inter-subject variability of functional responses and morphology in superior temporal cortex (Bonte et al., 2013). Second, we attempted to minimize the contribution of explicit top-down inferential processing, asking participants to focus on visual aspects of the scene rather than instructing subjects to make explicit social judgements.

In the present study, we asked if pre-adolescent children (aged 6 – 12 years) show *interaction-selective* responses (i.e. interaction > non-interaction) in the pSTS (pSTS-I), and if so, whether selectivity would differ from that of adults. We also explored responses in 4 other socially-tuned posterior temporal regions that might also plausibly show age-group differences in response to interactive stimuli: EBA, face-selective STS (STS-F), fusiform face area (FFA), and mentalizing-selective temporo-parietal junction (TPJ-M). Finally, we ran exploratory analyses to determine if further developmental trends existed, as evidenced by differences between sub-groups of younger children (6 – 8 year olds), older children (9 – 11 year olds), and adults.

## 2. Methods

### 2.1. Participants

31 children aged between 6 – 12 years (mean age = 8.94; SD = 1.88; 13 females) took part in the experiment, along with 29 adults (mean age = 23.14 years; SD = 4.21; range = 18 - 35; 16 females). All subjects were right hand dominant. Children gave informed assent (consent was also given by a guardian of each child) and they received gift vouchers (or toys of equivalent value) as compensation for participation. Adult subjects gave consent and received monetary compensation for participation. Ethical procedures were approved by the Bangor University psychology ethics board.

### 2.2. MRI Tasks & Experimental Session

All scans were acquired in one session that was split into two halves with a short break where subjects came out of the scanner for approximately 5 – 10 minutes. This served to minimize fatigue in children, but for consistency, adults also took this break. Additionally, children also completed a head-motion ‘training-session’ prior to entering the scanner (see supplementary materials A). Inside the scanner, two different video tasks were used to localize brain regions that are sensitive to: 1) Dynamic social interactions; and 2) dynamic faces and bodies. For both of these tasks, subjects were not instructed to make explicit judgements about the stimuli, but instead to simply to watch the videos.

The ‘social interaction localizer’ was almost identical to that used previously (Isik et al., 2017; Walbrin et al., 2018) and consisted of three runs of videos from three conditions: Interaction (i.e. two profile-view human point-light figures interacting with each other), non-interaction (i.e. two profile-view human point-light figures performing non-interactive actions, for example, one figure jumping, the other cycling), and scrambled interaction (i.e. average ‘motion-matched’ scrambled versions of the interactive stimuli where the coordinates of each point-light dot were randomly shifted to disrupt the perception of interactive or biological motion) (block length = 16s, based on three videos of variable length that summed to 16s; run length = 144s). Each run consisted of two blocks per condition – one presented in either half of each run – in randomized order with the other conditions.

Interaction-selective pSTS-I ROIs were localized with the interaction > scrambled interaction contrast. This not only captured differences in interactive content, but also biological motion (unlike the more ‘closely matched’ interaction > non-interaction contrast that does not capture large differences in biological motion). This ‘broader’ contrast was chosen as it was more comparable to other localizer contrasts that were used here, and are typically used elsewhere (e.g. Julian et al., 2012; faces > objects, rather than the relatively more ‘socially matched’ faces > bodies), and to account for the possibility that weaker interaction responses in children may have resulted in poorer localization of pSTS-I ROIs (i.e. children’s responses may have been weaker).

The face and body localizer was adapted from stimuli used previously (Pitcher et al., 2011), and served to localize face selective STS-F and FFA, along with body selective EBA. Participants completed three runs that contained blocks of videos that depicted either moving faces, moving bodies, or moving objects (STS-F & FFA localization contrast = faces > objects; EBA localization contrast = bodies > objects; block length = 18s (6 × 3s videos); run length = 270s). Each run consisted of four blocks per condition.

Additionally, mentalizing-selective temporo-parietal cortex (TPJ-M) was localized using the Pixar short-film ‘Partly Cloudy’ (2009), by modelling responses to timepoints that reliably evoke responses to mentalizing (along with ‘pain’, ‘social’ and ‘control’ time-points) as described previously (Richardson et al., 2018). However, as this region did not respond strongly to interaction stimuli, these data are not reported in the main text (see supplementary materials C for further details).

### 2.3. MRI Parameters, Pre-processing, & GLM Estimation

Scanning was performed with a Philips 3T scanner at Bangor University. The same fMRI parameters were used for all localizer tasks as follows: T2*-weighted gradient-echo single-shot EPI pulse sequence (with SofTone noise reduction); TR = 2000ms, TE = 30ms, flip angle = 83°, FOV(mm) = 240 × 240 × 112, acquisition matrix = 80 × 78 (reconstruction matrix = 80); 32 contiguous axial slices in ascending order, acquired voxel size (mm) = 3 × 3 × 3.5 (reconstructed voxel size = 3mm^3^). Four dummy scans were discarded prior to image acquisition for each run. Structural images were obtained with the following parameters: T1-weighted image acquisition using a gradient echo, multi-shot turbo field echo pulse sequence, with a five echo average; TR = 12ms, average TE = 3.4ms, in 1.7ms steps, total acquisition time = 136 seconds, FA = 8°, FOV = 240 × 240, acquisition matrix = 240 × 224 (reconstruction matrix = 240); 128 contiguous axial slices, acquired voxel size(mm) = 1.0 × 1.07 × 2.0 (reconstructed voxel size = 1mm^3^). Pre-processing (i.e. realignment & re-slicing, co-registration, segmentation, normalization, and smoothing) and general linear model (GLM) estimation were performed with SPM12 (fil.ion.ucl.ac.uk/spm/software/spm12). All SPM12 default pre-processing parameters were used except for the use of a 6mm FWHM Gaussian smoothing kernel, and all analyses were performed in normalised MNI space with 2mm isotropic voxels. Block durations and onsets for each experimental condition (per run) were modelled by convolving the corresponding box-car time-course with a canonical hemodynamic response function (without time or dispersion derivatives), with a high-pass filter of 128s and autoregressive AR(1) model. Head motion parameters (3 translation & 3 rotation axes) were modelled as nuisance regressors.

### 2.4. ROI Definition & Percent Signal Change Extraction

A group-constrained ROI definition procedure (e.g. Julian et al., 2012) was implemented as follows. Firstly, for each given localizer contrast (e.g. interaction > scrambled interaction), for each subject, a subject-specific ROI ‘search sphere’ was created by using *all other within-group subjects’* data to run a whole-brain analysis. This was used to find the voxel with the highest t-value for the corresponding localizer contrast around which an 8mm-radius search sphere was placed. For example, all adult group data, minus that individual, were used to create the search sphere for a given adult. This sphere size was chosen to ensure no overlap between search spheres for nearby but distinct regions (e.g. pSTS-I and STS-F) so that each ROI was comprised of unique voxels.

Subject-specific search spaces were then used to create the final set of ROIs for each participant, comprised of the 100 *most activated contiguous voxels* for the same contrast as that used to define the corresponding search space. The rationale for using 100 voxels was based on a compromise between using a much smaller number of voxels (e.g. 20 voxels) that might have resulted in exaggeratedly high selectivity values, and the maximum size of voxels within the search space (200+ voxels). However, 9 other similar sets of ROIs based on the highest 20, 40, 60, 80, 120, 140, 160, 180 and 200 voxels were also generated for replication and exploratory analyses.

For the interaction localizer as well as face and body localizer (for which there were 3 runs of data), a leave-one-run-out (LORO) approach was used to ensure that data used to define ROIs was independent of that used to extract percent signal change (PSC) values. Subject-wise PSC extraction for each condition within each left-out run was performed in MarsBaR (Brett et al., 2002) and the resulting values were averaged across all LORO iterations.

### 2.5. PSC & Selectivity Analyses

Mean PSC was extracted for each subject, for all tasks, yielding a total of 10 conditions: 3 interaction localizer conditions (interaction, non-interaction, & scrambled interaction) + 3 face and body localizer conditions (faces, bodies, & objects) + 4 mentalizing localizer conditions (mentalizing, pain, social, & control). PSC values were extracted from all 9 ROIs (4 bilateral ROIs and right FFA; left FFA could not be localized due to very weak responses across subjects). The main measure of interest – that is, *interaction selectivity* – was calculated with the following PSC subtraction: Interaction – non-interaction. For comparison analyses, face and body selectivity measures were also calculated as faces – objects, and bodies – objects, respectively. However, prior to these calculations, a series of one-sample t-tests were performed to determine which conditions activated each ROI above baseline. Above-zero PSC values for a given ‘target condition’ (i.e., interaction, faces, bodies, mentalizing) were considered a pre-requisite for calculating selectivity scores, as any region not univariately driven by a given target condition cannot not be interpreted as meaningfully responsive for that given category. Therefore, selectivity scores were not calculated for contrasts in regions that did not show above-zero PSC responses for given target conditions (see table 1).

**Table 1.**
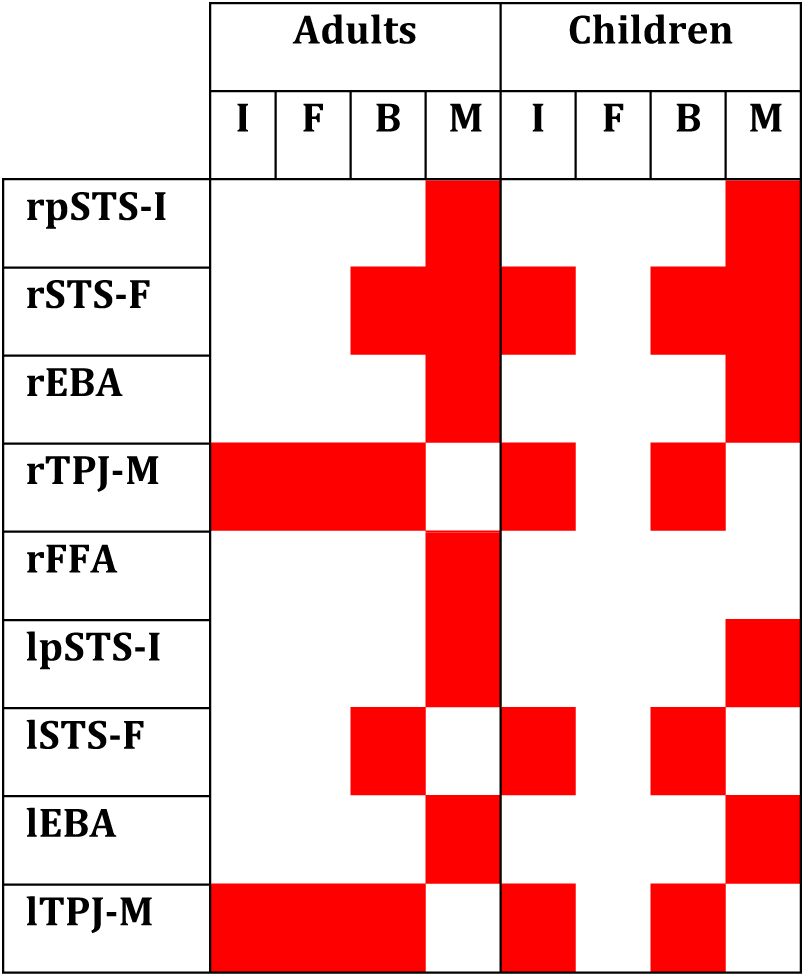
A table showing which selectivity measures were – and were not – calculated for each ROI. White cells denote significantly ‘above-zero’ PSC values for target conditions (red cells depict non-above zero PSC values). I = interaction; F = faces; B = bodies; M = mentalizing.

Finally, Bonferroni multiple-comparison correction was implemented for each ‘set’ of analyses separately (but not for exploratory analyses); the corrected Bonferroni threshold (α) is stated for each series of tests in the results section. All significant tests survived Bonferroni correction, unless otherwise stated. All one-sample t-test p*-*values are one-tailed.

### 2.6. Controlling for Head Motion

In line with previous developmental studies (e.g. Peelen et al., 2009), we excluded runs of data with > 2mm scan-to-scan movement across runs. This resulted in the exclusion of single runs of data in three separate children (although including these runs did not meaningfully change any results). In addition to removing these runs, differences in head motion between groups were tested using an analysis similar to Kang et al. (2003). Crucially, these results revealed that any age-group differences in selectivity are not attributable to head motion differences (see supplementary materials B for results of this analysis).

## 3. Results

### 3.1. Initial PSC Analyses

Before addressing the primary hypotheses, a 2×2 mixed ANOVA (condition x age-group) was performed with PSC values for each pSTS-I ROI, separately (Bonferroni corrected α = .025) to test for group differences between the interaction and non-interaction conditions. For the right pSTS-I (see figure 1), greater responses were observed in the interaction compared to non-interaction condition (*F*(1,58) = 34.27, *p* < .001). No main effect of age-group (*F*(1,58) = 0.18, *p* = .671) was observed, and the interaction between factors was marginally significant (*F*(1,58) = 2.83, *p* = .098) showing a small to medium effect size (*η*_*p*_^2^ = .046). This demonstrated that children’s responses to interaction conditions in the right pSTS-I are relatively adult-like, although a trend towards group differences exists.

**Figure 1.**
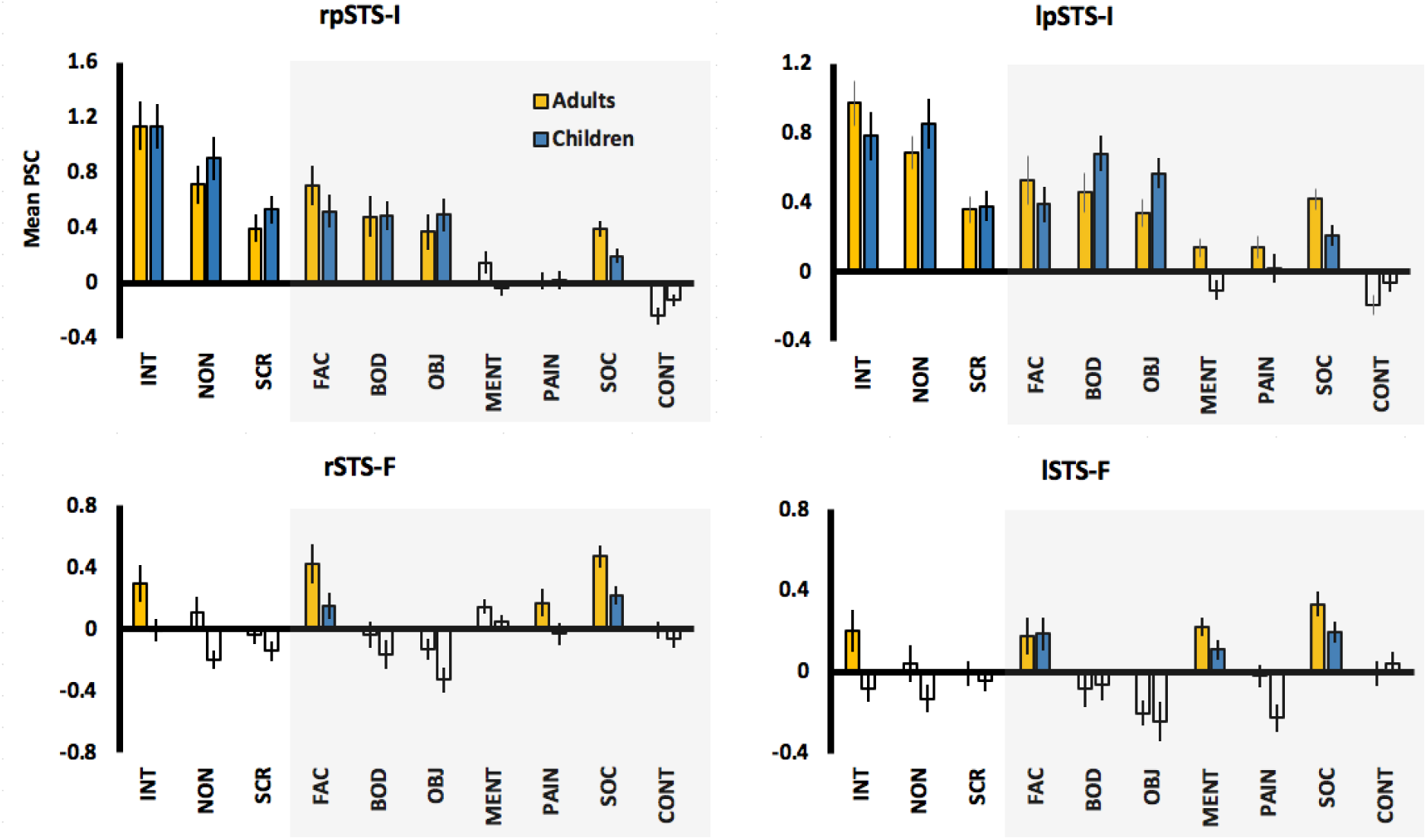
Mean PSC for the 3 interaction localizer conditions (unshaded region) along with other conditions (shaded region) in the right and left pSTS-I ROIs. Bilateral STS-F ROIs are also shown for comparison. White bars correspond to PSC values that were not significantly greater than zero (i.e. one-sample *p* value > .05). Error bars are SEM. INT = interaction; NON = non-interaction; SCR = scrambled interaction; FAC = faces; BOD = bodies; OBJ = objects; MENT = mentalizing; PAIN = pain; SOC = social; CONT = control.

In the left pSTS-I, a similar pattern of results was observed – a main effect of condition (*F*(1,58) = 4.47, *p* = .039; although this did not survive Bonferroni correction), no main effect of group (*F*(1,58) = 0.01, *p* = .944), but a significant interaction was shown (*F*(1,58) = 12.27, *p* = .001). This strong interaction reflects a clear difference between conditions in adults, but not in children (see table 2 & figure 3).

**Table 2.**
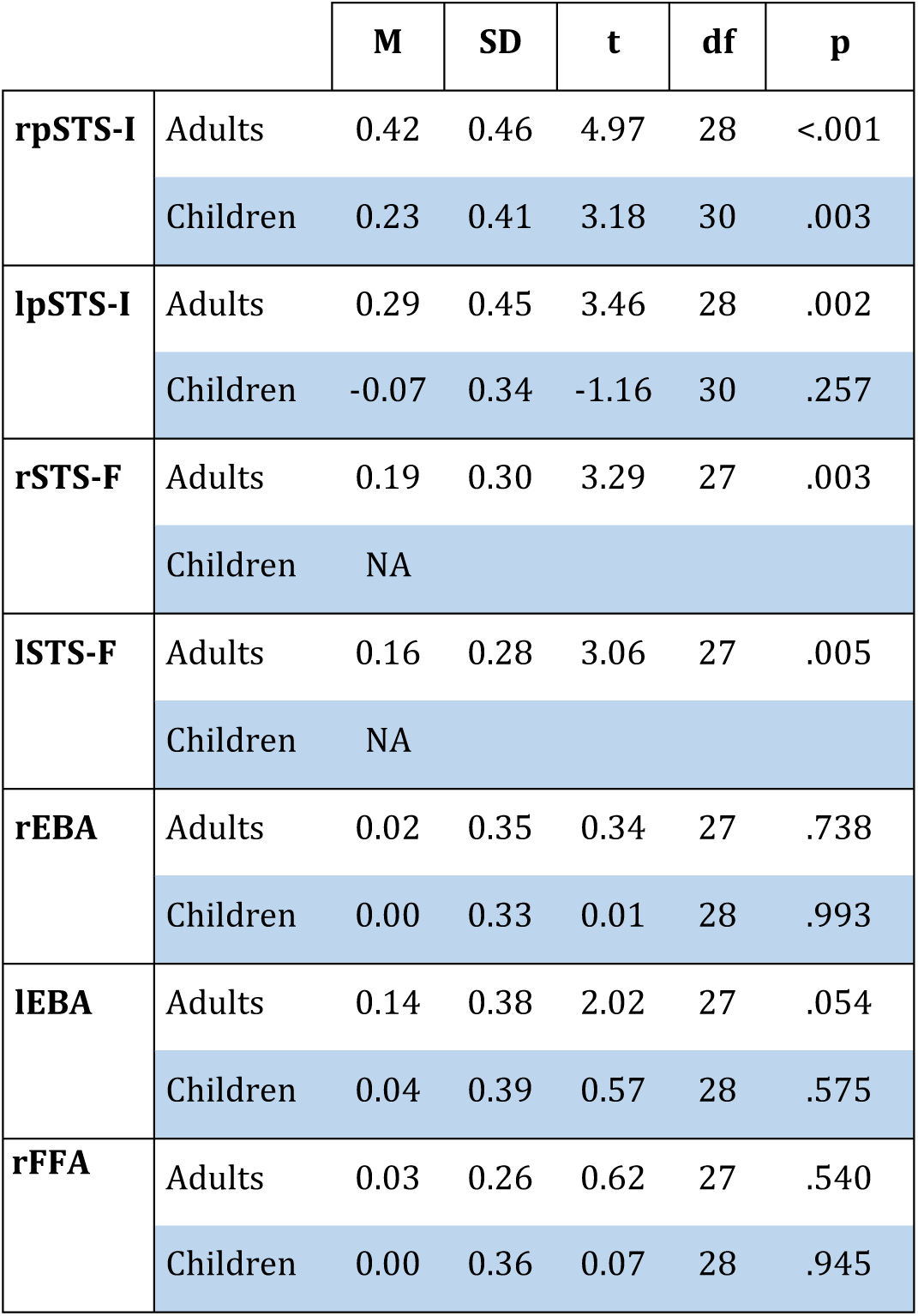
One-sample descriptive statistics for interaction selectivity measures in the bilateral pSTS-I, STS-F, EBA, and right FFA. NA = selectivity not calculated (i.e. interaction PSC was not greater than zero).

Informal inspection of PSC values in both pSTS-I ROIs showed stronger responses to the interaction condition than face, body, or mentalizing conditions, and also that responses were stronger for the interaction condition in the right than left pSTS-I, for both groups (*p* = .017; see supplementary materials D for full statistics). Additionally, no other region showed strong evidence for differentiation between interaction and non-interaction conditions except for bilateral STS-F ROIs in adults (see figure 1; see supplementary materials E for PSC charts for other ROIs). These findings also show that PSC responses in children are not uniformly weaker than adults; indeed, children showed approximately equivalent or greater PSC values than adults for several conditions across ROIs. Therefore, any differences in selectivity are not attributable to weaker overall responses in children.

### 3.2. Interaction Selectivity in pSTS-I

A series of analyses were performed to test interaction selectivity in each ROI (see figure 2). Regions for which selectivity was not calculated (i.e. due to non-above-zero interaction PSC values) are shown in table 1. One-sample t-tests (uncorrected) were performed to determine which regions showed above-zero selectivity for interactions (see table 2 for one-sample statistics for interaction selectivity values). For adults, strong interaction selectivity was shown for the right (*p* < .001) and left pSTS-I (*p* = .002), and bilateral STS-F (both *ps* < .005), and marginal selectivity in the left EBA (*p* = .054). By contrast, the right pSTS-I was the only interaction selective region in children (*p* = .002).

**Figure 2.**
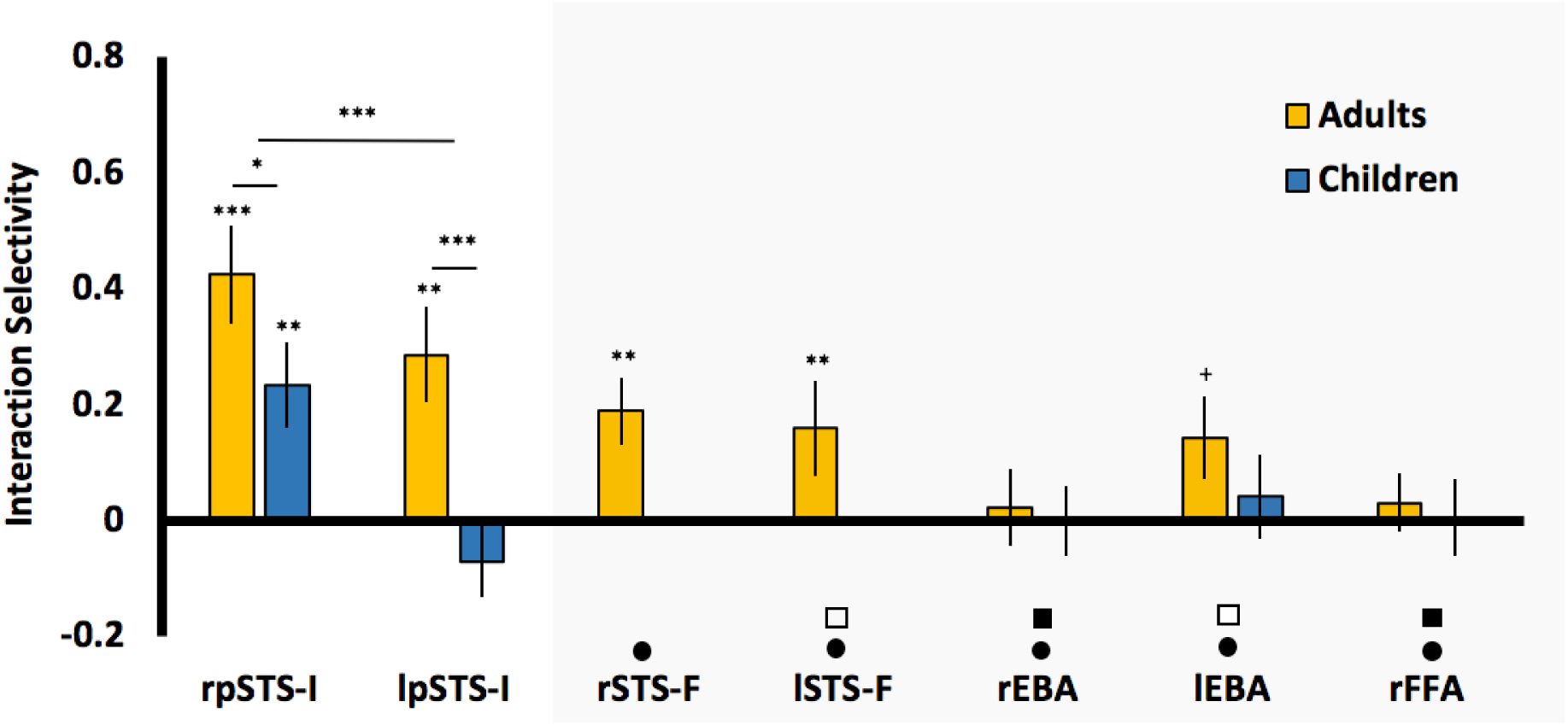
Mean interaction selectivity values for the pSTS-I ROIs (unshaded region) and other ROIs (shaded region). *** = *p* < .001; ** = *p* < .01; *= *p* = .049; += *p* = .054. Error bars are SEM. Black circles denote regions that are significantly less selective than right pSTS-I for both age-groups. Black and white squares, respectively, denote regions that are either significantly or marginally less selective than the left pSTS-I in adults.

To test the hypothesis that adults are more interaction selective than children in the pSTS-I, two independent t-tests were performed (Bonferroni corrected α = .025) on selectivity values in right and left pSTS-I, separately (due to a main effect of hemisphere; see supplementary materials F). In line with the interaction terms in the initial PSC ANOVAs, adults were significantly more selective than children in the left hemisphere (*t*(58) = 3.50, *p* = .001). This effect was also observed in the right hemisphere, but did not survive multiple comparison correction (*t*(58) = 1.68, *p* = .049). These findings show greater interaction selectivity for adults than children in the left pSTS-I, as supported by a large effect size (Cohen’s *d* = 0.90), with a smaller difference in the right pSTS-I, as supported by a small-to-medium effect size (Cohen’s *d* = 0.44). Although both groups show selective responses in the right pSTS-I, unlike adults, children do not show interaction selectivity in the left pSTS-I.

### 3.3. pSTS-I is More Interaction Selective than Other ROIs

A series of analyses were then performed to assess whether interaction selectivity was significantly greater in the pSTS-I than other ROIs (see supplementary materials F for full statistics; see figure 2). These results showed that for both adults and children alike, selectivity was significantly greater in the right pSTS-I than all other ROIs (all *ps* < .032). For the left pSTS-I, only adults showed above-zero interaction selectivity, and so analyses were only performed with adult data. Statistically significant results were shown (at either a corrected or uncorrected threshold) for comparisons against bilateral EBA, right FFA and left STS-F (all *ps* < .044); however, the trend towards greater selectivity in the left pSTS-I compared to right STS-F was not significant (*p* < .113).

### 3.4. Interaction vs. Face & Body Selectivity

Interaction selectivity is greater in the pSTS-I ROIs than virtually all other ROIs, but is the pSTS-I more selective for interactions than for faces or bodies? To test this, a 3 × 2 mixed ANOVA (selectivity category x age-group) with follow-up tests was performed for the right pSTS-I, along with t-tests in adults for the left pSTS-I (Bonferroni corrected α = .01; see supplementary materials G & H for charts and full statistics of face and body selectivity across ROIs). In the right pSTS-I, a main effect of selectivity category (*F*(1.46,80.45) = 8.11, *p* = .001), and a main effect of age-group at an uncorrected level was shown (*F*(1,55) = 4.92, *p* = .031). As the interaction term was not significant (*F*(1.46,80.45) = 1.05, *p* = .337), follow-up paired t-tests were performed on both adults and children’s selectivity scores together. Interaction selectivity was significantly greater than body selectivity (*t*(56) = 4.17, *p* < .001), and greater than face selectivity at an uncorrected threshold (*t*(56) = 1.75, *p* = .043).

For the left pSTS-I, adults showed greater interaction selectivity than body selectivity at an uncorrected threshold (*t*(27) = 2.12, *p* < .022), and the trend for greater interaction than face selectivity was not significant (*t*(27) = 1.12, *p* = .132). These results demonstrate that, for both adults and children alike, interaction selectivity in the right pSTS-I is greater than body selectivity, and to a lesser extent, greater than face selectivity. These effects are considerably weaker for adults in the left pSTS-I (and absent in children).

### 3.5. Whole-brain Analyses

To determine whether other regions outside of the functionally localized ROIs demonstrated sensitivity for the interaction > non-interaction contrast, whole-brain analyses were performed for adults and children. The resulting group-level t-maps were height-thresholded at *p* = .001, and false discovery rate (FDR) cluster-corrected at *p* < .05. For adults, right hemisphere responses were shown with peak activation in the pSTS, along with activations extending to the anterior STS (aSTS), and a small cluster in IFG. Additionally, bilateral precuneus and small left calcarine sulcus activation was shown. For children, similar although weaker responses were shown in the right pSTS and aSTS regions only. When comparing activation between the two groups directly, no differences were observed for either the adults > children, or the reverse contrast, demonstrating that these analyses were not as sensitive as ROI analyses at capturing group differences.

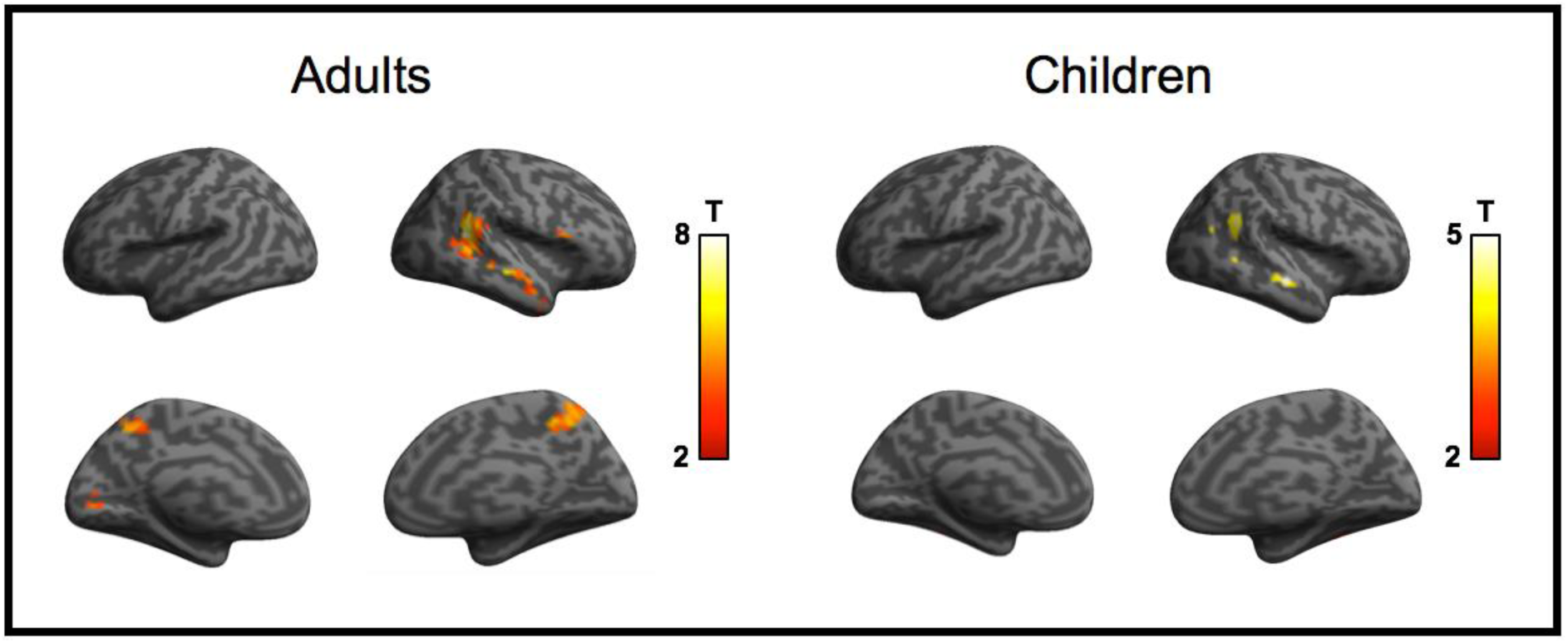
Whole-brain activation for the interaction > non-interaction contrast for adults and children. Colour bar represents activation t-value. Height threshold = .001; FDR cluster correction = .05.

### 3.6. Exploratory Analysis: pSTS-I Interaction Selectivity as a Function of ROI Size

The preceding analyses used selectivity measures that were generated from ROIs with a fixed size of 100 voxels, therefore ensuring no group differences in ROI size. However, without knowing the underlying spatial organization of selective responses within the pSTS – and crucially, whether this differed between groups – it is possible that the current choice of ROI size might have favoured adult responses. For example, selecting the highest 100 contiguous voxels in adults might approximately capture the entire peak of a highly interaction selective cluster, whereas this peak region might be smaller in children, and as such, selectivity measures in children may have been calculated with the inclusion of less selective voxels outside of this hypothesized peak. Exploratory analyses were conducted to determine if selectivity differed as a function of ROI size, and whether such changes were similar between groups.

Interaction selectivity as a function of ROI size is shown in figure 4. A clear trend for greater selectivity in adults than children is shown across all ROI sizes. Interestingly, adults show an approximately linear decrease in selectivity with increasing ROI size, whereas this does not appear to be the case for children. To formally test this apparent trend, linear regression slopes were calculated for each subject; that is, a beta coefficient that describes the change in selectivity across ROI sizes was calculated, per ROI, per subject, and entered into a series of (uncorrected) group tests (see supplementary materials I). These analyses revealed that beta coefficients were significantly more negative for adults than children in bilateral pSTS-I (*p* = .010); these findings show that interaction selectivity decreases linearly with increasing ROI size in adults, but not children. We speculate that this might indirectly reflect age-related differences in the *focal tuning* of interaction selective responses in the pSTS; that is, adults show strongest selectivity around a small peak cluster in the pSTS with intermediate selectivity in neighbouring STS cortex, while children are less selective overall and do not show graded changes in selectivity in the right pSTS (and are not selective at all in the left pSTS).

**Figure 4.**
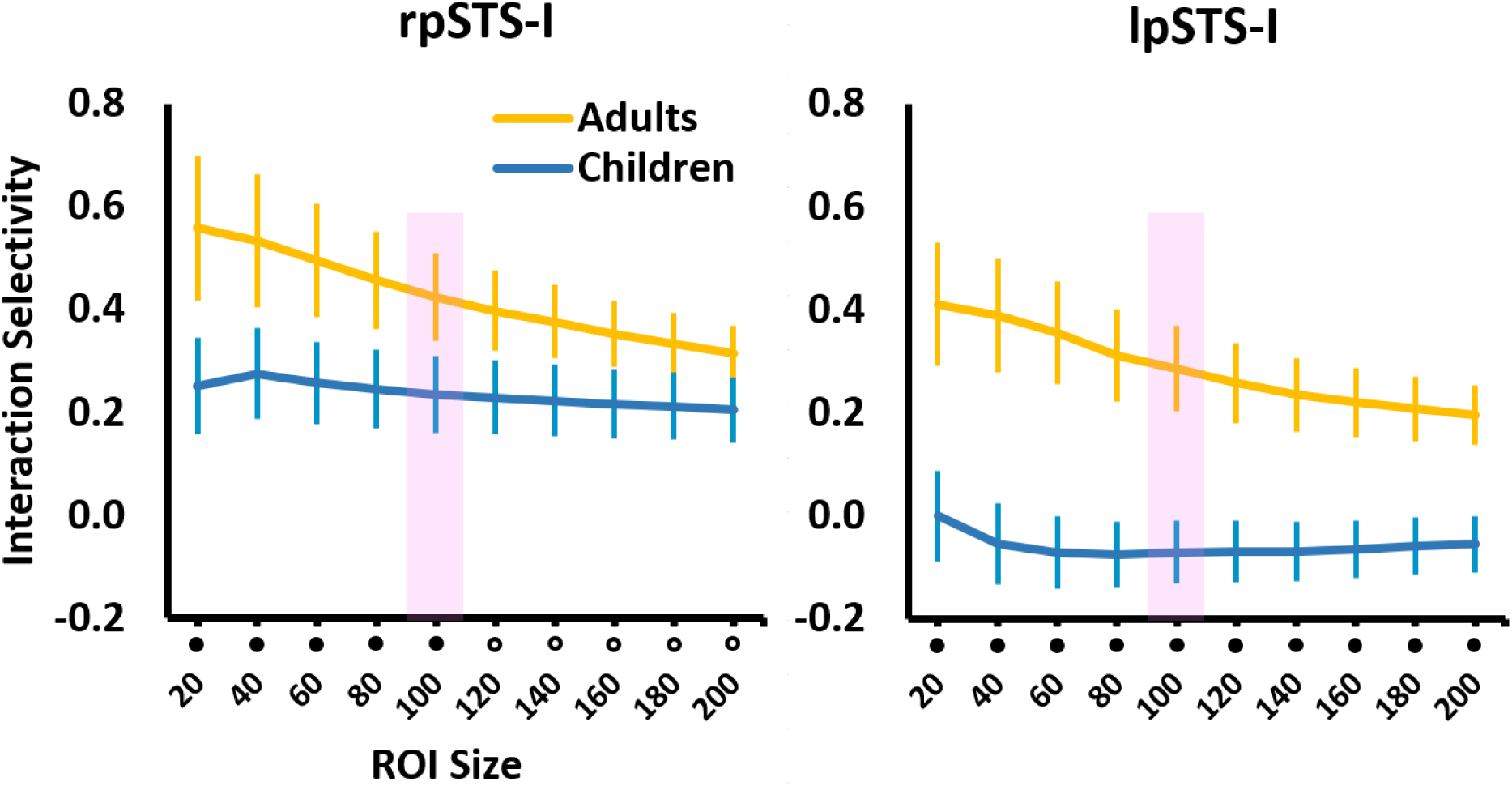
Mean interaction selectivity plotted as a function of ROI size, for both age-groups, for the right and left pSTS-I. Black filled circles denote ROI sizes where selectivity was significantly greater for adults than children (*p* < .05). Unfilled circles denote marginally greater selectivity for adults than children (*p* < .10). The shaded region shows the 100 voxel ROI, as used in the preceding analyses. Error bars are SEM.

### 3.7. Exploratory Sub-group Analyses: Interaction Selectivity

The current study was designed to compare differences between adults and pre-adolescent children (aged 6 – 12 years), but we are also interested in whether there is developmental change *across age* within the child group. To test if this might be true, we compared interaction selective responses in pSTS-I across smaller sub-groups: 6-8 years (N = 14), 9-11 years (N = 14), and a random subset of adults (N = 14). Despite the relatively small group sizes, this exploratory analysis was intended to see whether nuanced sub-group differences in interaction selectivity might exist.

Sub-group differences in interaction selectivity were observed in the right pSTS-I for the 100 voxel ROI (see figure 5; see supplementary section J for full statistics). Adults were significantly more selective for interactions than 6-8 year olds (*p* = .035) but not 9-11 year olds (*p* = .961), while 9-11 year olds were more selective than 6-8 year olds (*p* = .044). For the left pSTS-I, interaction selectivity was significantly greater for adults than both 6-8 year olds (*p* = .001) and 9-11 year olds (*p* = .035), and a marginal trend for greater selectivity for 9-11 year olds than 6-8 year olds was also observed (*p* = .063). These findings suggest that the marginal trend observed in the right pSTS-I in the main interaction selectivity analysis (i.e. all adults > all children) is likely driven by weaker selectivity in younger children; by contrast, both older and younger children are less selective than adults in left pSTS-I, and selectivity appears to increase with age. Importantly, although these results were obtained with the 100 voxel ROIs, they are robust across different ROI sizes.

**Figure 5.**
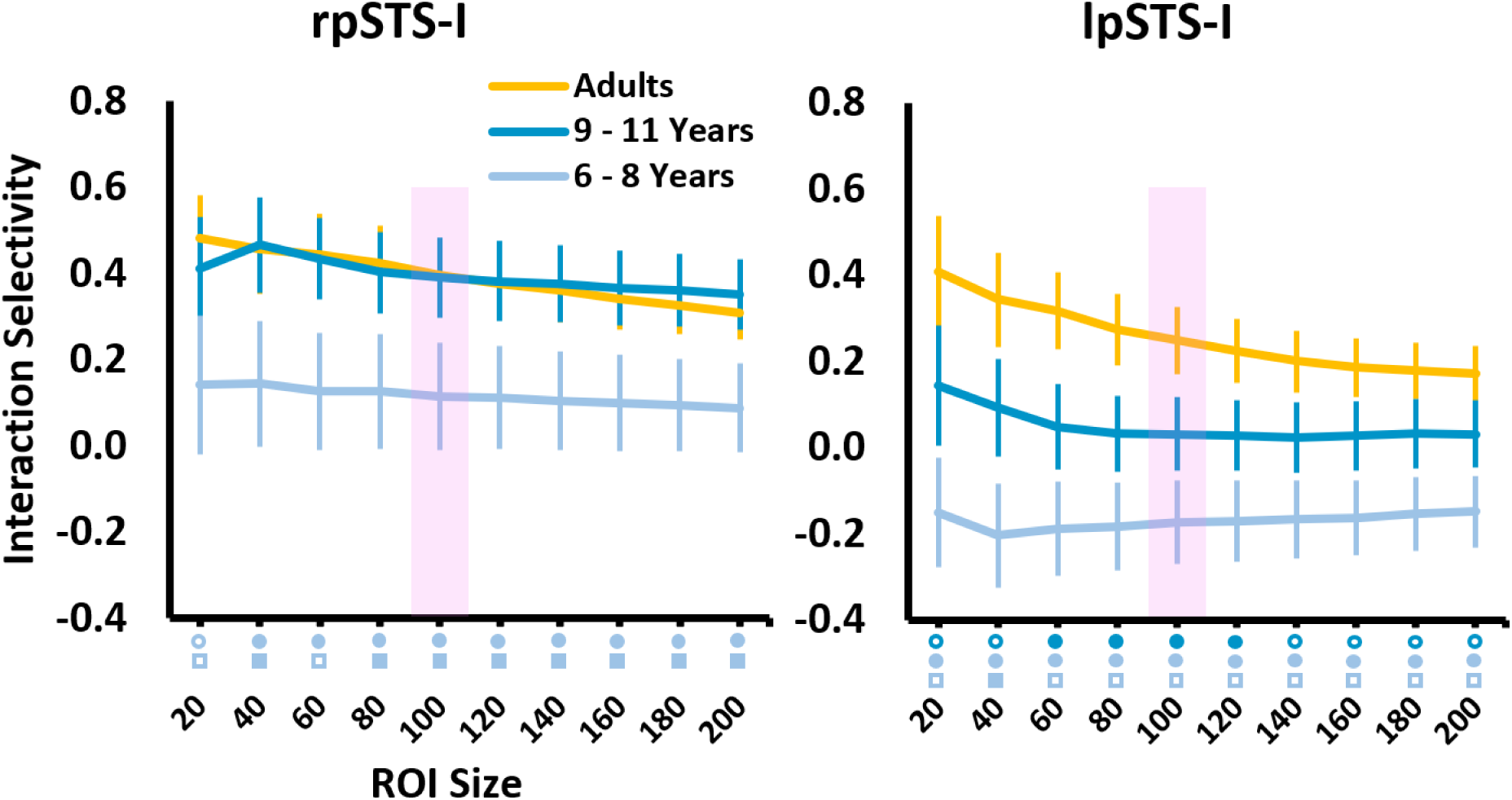
Mean interaction selectivity plotted as a function of ROI size, for the three sub-groups for the right and left pSTS-I. The shaded region denotes the original 100 voxel ROI. For each ROI size, statistical effects for each sub-group t-test comparison are indicated by coloured symbols underneath the x-axis, as follows; dark blue circles: adults > 9 –11 years; light blue circles: adults > 6 – 8 years; light blue squares: 9 – 11 years > 6 – 8 years; filled symbols: *p* < .05; unfilled symbols: *p* < .10. Error bars are SEM.

As with the all adults > all children comparison, ROI size regression slope analyses were conducted to determine if group differences in these negative linear trends were observed between the three sub-groups; these analyses demonstrated a linear decrease in selectivity with ROI size for bilateral pSTS-I for adults, but not for either group of children (see supplementary materials K for statistics). Additionally, whole-brain analyses in the three sub-groups show right pSTS activation for the interaction > non-interaction contrast in adults and older children, but not younger children, along with greater pSTS responses for adults compared to younger, but not older children (see supplementary materials L for full results).

## 4. Discussion

Here, we report four key findings. First, and importantly, children as young as six show selective responses to social interactions in the pSTS, implying that even in childhood, the social brain is already tuned to understand social interactions. Secondly, as predicted, adults showed greater selectivity for social interactions than children in the pSTS; this effect was strongest in the left hemisphere, where children showed no selectivity, but was only marginal in the right hemisphere, where children showed intermediate levels of interaction selectivity. Exploratory analyses further elaborated these trends and showed that the all adult > all children difference in the right pSTS is likely driven by weaker selectivity in the youngest children (6 – 8 years), while older children (9-11 years) already demonstrated adult-like selectivity. Developmental trends in the left pSTS were intuitive, showing a graded pattern of selectivity across the three sub-groups (i.e. adults > older children > younger children). Importantly, both the main and sub-group findings generalized across different ROI sizes, suggesting that these are stable effects. Thirdly, and unexpectedly, unlike children, adults showed additional interaction-specific co-activations in other ‘socially tuned’ temporal lobe regions. Fourthly, adults demonstrated much more focally tuned interaction responses in bilateral pSTS, where selectivity dropped off from a core ‘peak’, whereas children showed weaker, broader selectivity that did not change across ROI size. Altogether, although children do already show functional selectivity for social interactions, these findings demonstrate that neural responses to social interactions are not fully mature in pre-adolescent children, and therefore must undergo substantial development during adolescence.

Both adults and children showed significantly greater interaction selectivity in the right pSTS than all other ROIs, and this was supported by the whole-brain findings that show this was the most active region. These findings are consistent with previous accounts that the pSTS – especially in the right hemisphere – is strongly responsive to dyadic social interactions (Georgescu et al., 2014; Isik et al., 2017; Kujala, Carlson, & Hari, 2012a; Lahnakoski et al., 2012; Walbrin et al., 2018; Walbrin & Koldewyn, 2019).

Adults showed stronger interaction selective responses in the pSTS than children, especially in the left-hemisphere, in line with analogous trends for functionally localized face (Scherf et al., 2007) and body regions in the STS (Ross et al., 2014). Additionally, adults showed strong responses in both hemispheres, and greater responses in right than left pSTS. By contrast, children demonstrated above-zero selectivity in the right but not left pSTS. Interestingly, similar developmental changes have been observed previously in the STS, albeit in a different modality. Bonte et al. (2013) calculated laterality scores for voice selectivity in the STS (i.e. the ratio between the magnitude and extent of left and right lateralized STS responses) and found that both children and adults showed rightward lateralization. Crucially, this effect was significantly stronger in children, as proportionally more recruitment of left STS was shown for adults. Additionally, voice selective responses in the STS were more diffuse and less selective in children than adults (who showed both strong and spatially constrained selectivity), mirroring age related differences in tuning of interaction selectivity in the current study. These results suggest that selective responses in the STS may become relatively more bilateral and focally tuned across development (although this trend is not pronounced for faces in our data; see supplementary section M).

It is also worth noting that ‘non-selectivity’ for interactions in the left pSTS in children is not the result of weaker interaction responses per se; instead, strong PSC responses were found for both interactions and non-interactions, but they were approximately equal in magnitude. This could be interpreted in two ways. Firstly, that responses in this region are driven by the mere presence of two individuals, but not by interactive information (e.g. contingent actions and facing direction). As such, immature responses in this region may reflect simplistic representations of interactions as merely two people together, that are insensitive to nuanced dynamic information that adults make use of to distinguish these two conditions. And secondly, responses in this region may simply reflect sensitivity to biological motion per se, that was essentially equivalent between interaction and non-interaction stimuli. This is supported by previous evidence of STS sensitivity to biological motion in children (e.g. Mosconi et al., 2005; Carter & Pelphrey, 2006) and additionally, PSC responses to the body localizer task were comparable to interactions in the left pSTS but not right pSTS. Therefore, it seems that left pSTS responses to interaction stimuli are specific to general biological motion information in children, and become more sensitive to interactive information across development.

Further to this, adults, but not children, unexpectedly showed complementary interaction selective responses in regions neighbouring interaction selective pSTS (e.g. face selective STS cortex and to a marginal extent, left EBA). Along with strong focal tuning in the pSTS, weaker selectivity in neighbouring face selective STS is somewhat unsurprising, given the relevance of facial information in interactive contexts. Indeed, a similar pattern was shown for face information, whereby stronger face selective responses were found in face selective STS, with weaker face selectivity in neighbouring interaction selective pSTS in adults (see supplementary materials G). Partially overlapping STS responses are shown for other categories of social information (e.g. theory of mind, biological motion, faces, and voices; Deen et al., 2015; Lahnakoski et al., 2012) that are likely important to understanding the content of social interactions. While previous research in adults has demonstrated pSTS-I responds selectively even when interactions do not contain any human information (i.e., when interactants are animate moving shapes; Isik et al., 2917; Walbrin et al., 2018), our data suggests that other social regions in the temporal lobe may also, by adulthood, develop at least some sensitivity to social interaction cues. While the present findings focus on dynamic whole-body dyadic interactions, further research could address the possibility that such ‘interaction tuning’ varies by social category along the extent of the STS (e.g. posterior STS interaction selectivity for bodies may be complemented with more anterior tuning to face- and voice-based interactions).

The whole-brain findings reported here are somewhat similar to the results of Sapey-Triomphe et al. (2017; i.e. using a highly similar stimulus set and the same interaction > non-interaction contrast). In this study, the authors reported strong pSTS (and wider posterior temporal cortex) responses across adults, adolescents, and children (8-11 years), along with activation in IFG. These results are largely replicated by the present analyses; both adults and children (but not younger children in sub-group analyses) showed responses in right pSTS and aSTS. Although we did not predict responses in the aSTS, co-activation of pSTS and aSTS have been previously shown for social interactions (Lahnakoski et al., 2012), and dynamic faces (Pitcher et al., 2011), and may suggest functional coupling between these regions, although whether these regions perform similar or different computations during interaction perception remains to be seen.

The present findings demonstrate that the right pSTS in pre-adolescent children already shows selectivity to social interactions, similar to that seen in adults. However, children also show markedly different neural responses to dyadic social interactions. The present results imply that the maturation of neural interaction selectivity is characterized by increasingly bilateral pSTS activation, increasingly stronger selectivity and more focal tuning bilaterally, and increased, though non-selective, responses to social interaction in neighbouring temporal cortex. These findings motivate further research to explore interaction responses across the full life span, including adolescence and late adulthood, and to explore developmental changes in the context of other types of interactions (e.g. stimuli that are restricted to faces or voices).

