## Supplementary Materials for "Developmental Changes in Visual Responses to Social Interactions"

### *A. Head Motion Training Session*

Prior to entering the scanner, children (but not adults) completed a short head motion 'training session'. This entailed lying still inside a 'mock scanner' with a motion sensitive electrode (https://pstnet.com/products/motrak/; MoTrak Head Motion Tracking System) placed on the forehead to measure movement across three translation and three rotation axes. Children viewed a monitor through a mirror attached to an MRI head coil, allowing for visual motion feedback (i.e. an on-screen cursor that was 'controlled' by head movements was visible to subjects, as well as researchers via a separate monitor). Pre-recorded audio of an fMRI acquisition sequence was also played during this session to further simulate the real scanner environment.

Subjects were instructed to lie still and 'keep the cursor in the middle of a target circle' that was presented centrally on screen. The diameter of this circle corresponded to a region allowing for 3mm head translations in any direction. Once participants were able to keep the cursor within this target region for a timed period of 30 seconds, a different task was performed without visual motion feedback; subjects were simply instructed to stay relaxed and keep still while watching a short animated video but were told that if they moved 'too much' (i.e. > 3mm translation movements from a their initial head position), video playback would be paused, signalling that they had moved too much. Once they were able to watch the video for a period of 2 minutes without the video pausing, they were deemed 'ready' to be scanned. Head movement data was not recorded or analysed, as the sole purpose of this session was to train children to remain still inside the scanner.

### *B. Head Motion Analysis*

A head motion analysis was performed to determine whether potential group differences in selectivity were attributable to head motion differences. To test this possibility, a similar approach to Kang et al. (2003) was adopted. That is, a root mean square (RMS) measure of head motion across the six motion parameters was calculated across all time points *for each localizer condition separately* (e.g. for interaction and non-interaction, separately) per run, for each subject. These head motion values were averaged across runs to form a single measure per condition, for each subject, that was entered into ANOVAs and t-tests.

For the interaction selectivity contrast (i.e. interaction vs. non-interaction RMS head motion values), a 2 x 2 mixed ANOVA (condition x age-group) was performed. Crucially, this did not reveal a main effect of condition ($F(1,58) = 0.00$, $p$ =.968). A significant main effect of group ($F(1,58) = 13.50$, $p$ =.001) but no interaction between factors was also shown ($F(1,58) = 0.31$, $p$ =.580). Analogous findings were shown for face selectivity (i.e. faces vs. objects): A main effect of group ($F(1,56) = 12.66$, $p$ =.001) but neither main effect of condition nor interaction term reached significance (both $ps$ > .696). Similarly, for body selectivity (i.e. bodies vs. objects), a main effect of group ($F(1,56) = 12.24$, $p$ =.001, but neither main effect of condition nor interaction reached significance (both $ps$ > .415). Therefore, despite greater head movement overall for children, head motion cannot account for any selectivity differences for these contrasts, as head motion was matched for each pair of conditions used to calculate selectivity measures.

### C. *Mentalizing Localizer*

To localize mentalizing selective temporo-parietal cortex (TPJ-M), participants viewed the Pixar short-film ‘Partly Cloudy’ (2009; duration = 355s, including 10s rest). Previous research (Richardson et al., 2018) used a reverse-correlation analysis to identify time points within the video that reliably evoke responses to mentalizing, along with ‘pain’, ‘social’, and ‘control’ time-points, allowing for the localization of mentalizing and pain network regions (total time per condition: Mentalizing = 44s; pain = 26s ; social = 28s; control = 24s). Bilateral TPJ-M ROIs were localized by contrasting responses during mentalizing timepoints with pain timepoints (i.e. mentalizing > pain).

For definition of TPJ-M and extraction of mentalizing localizer responses, a similar approach to other ROIs/tasks was implemented, except that given only one run of mentalizing data, extraction and definition were not independent of each other in the TPJ-M; however, mentalizing responses were not intended to be measured in this region and extraction of interaction and face & body localizer conditions remained independent. It is important to note that for each run of data for the interaction and face & body localizer tasks, responses were extracted and averaged, in an identical manner as for other ROIs (see main text).

Importantly, TPJ-M ROIs were excluded from selectivity analyses (i.e. interaction, face, & body selectivity) as the mentalizing condition was the only target condition to show above-zero responses in both groups in these regions (although children showed some sensitivity to faces in these regions; see section E, below).

*D. Initial PSC Analyses: Hemisphere x Condition pSTS-I ANOVA*

To determine whether PSC for the interaction condition was greater in the right than left pSTS-I, a 2x2 mixed ANOVA (hemisphere x age-group) was performed. This revealed a main effect of hemisphere ($F(1,58) = 6.05$, $p = .017$), but no main effect of group ($F(1,58) = 0.27$, $p = .607$), or interaction between factors ($F(1,58) = 0.81$, $p = .373$), demonstrating stronger responses in the right pSTS-I in both groups.

*E. PSC Charts for Bilateral EBA, TPJ-M, & Right FFA*

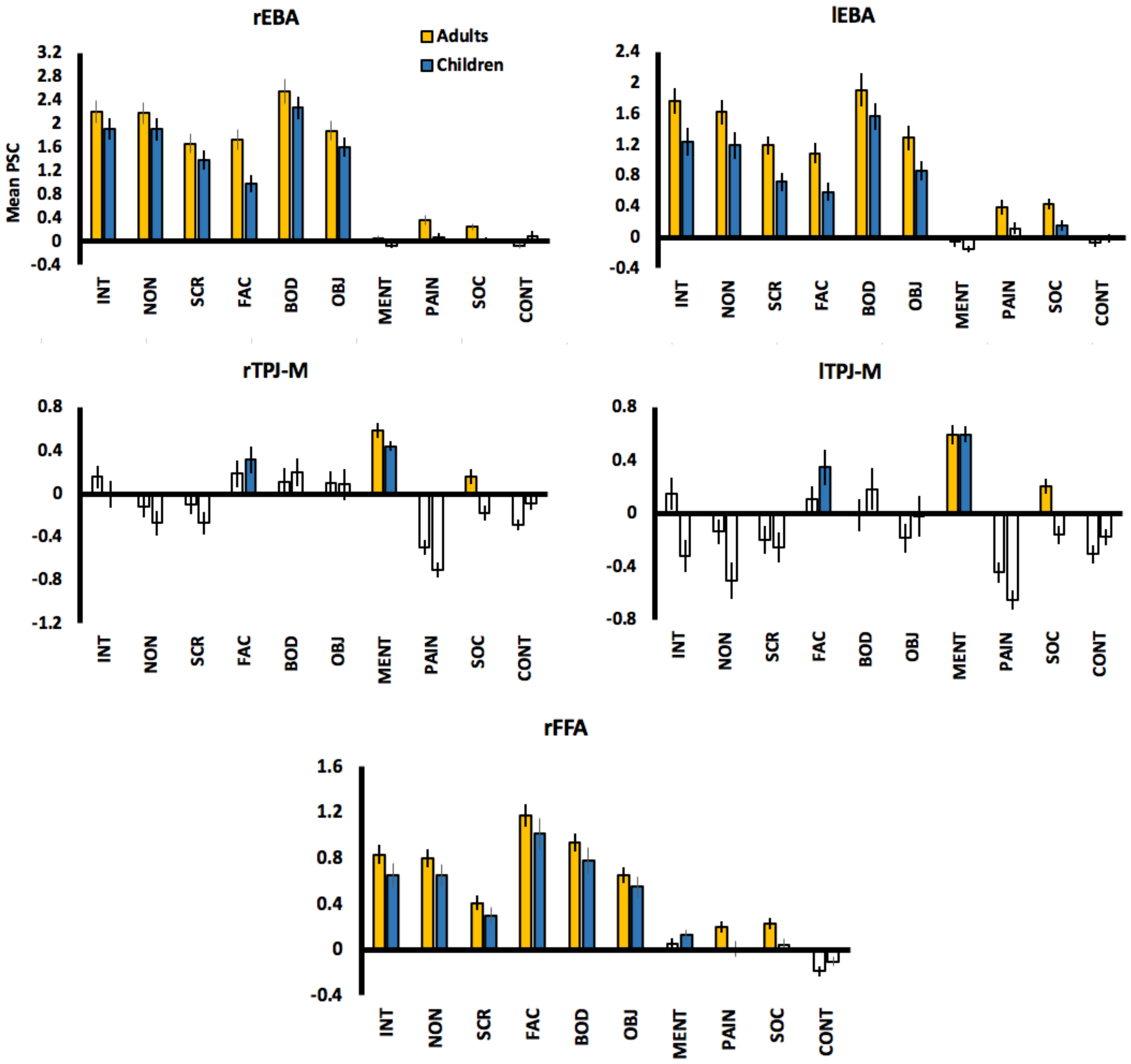


Mean PSC for the 3 interaction localizer conditions (unshaded region) along with other conditions (shaded region). White bars correspond to PSC values that were not significantly greater than zero (i.e. one-sample *p* value > .05). Error bars are SEM. INT = interaction; NON = non-interaction; SCR = scrambled interaction; FAC = faces; BOD = bodies; OBJ = objects; MENT = mentalizing; PAIN = pain; SOC = social; CONT = control.

*F. pSTS-I vs. Other ROIs: Interaction Selectivity*

A series of ANOVAs and t-tests were used to test whether interaction selectivity is greater in the pSTS-I than other ROIs (Bonferroni corrected $\alpha = .006$). First, right pSTS-I and left pSTS-I were compared with a 2 x 2 mixed ANOVA (hemisphere x age-group). Greater responses were observed in the right pSTS-I ($F(1,58) = 21.70$, $p < .001$), and greater selectivity overall was shown for adults ($F(1,58) = 8.05$, $p = .006$), along with a marginal interaction, suggesting a trend for greater selectivity in adults compared to children ($F(1,58) = 3.11$, $p = .083$). Follow-up paired t-tests showed that the trend for greater right than left pSTS-I selectivity was statistically stronger in children ($t(30) = 4.83$, $p < .001$) than adults ($t(28) = 1.93$, $p = .032$; significant at uncorrected level only). These findings demonstrate strongly right lateralized interaction selectivity in children, while adults show more bilateral pSTS-I selectivity that is slightly stronger in the right hemisphere.

To determine if right pSTS-I selectivity was greater than for other regions in which *both groups* showed above-zero selectivity (i.e. right EBA, left EBA, & right FFA), 2 x 2 mixed ANOVAs (region x age-group) were performed. When compared to the right EBA, a main effect of region ($F(1,55) = 36.09$, $p < .001$), no main effect of group ($F(1,55) = 1.78$, $p = .187$), and marginal interaction ($F(1,55) = 3.18$, $p = .080$) was shown. For the left EBA, a main effect of region ($F(1,55) = 16.39$, $p < .001$), no main effect of group ($F(1,55) = 2.98$, $p = .090$), and no interaction ($F(1,55) = 0.86$, $p = .357$) was observed. And for the right FFA, a main effect of region ($F(1,55) = 36.43$, $p < .001$), no main effect of group ($F(1,55) = 1.96$, $p = .168$), and marginal interaction ($F(1,55) = 3.19$, $p = .080$). Therefore, right pSTS-I selectivity was greater than all three comparison regions, with no main effect of group. Marginal interaction trends for the right EBA and right FFA comparison were shown, and so follow-up paired t-tests (one-tailed) were conducted and revealed the following significant effects: Right pSTS-I > right EBA in adults ($t(27) = 4.71$, $p < .001$); right pSTS-I > right EBA in children ($t(28) = 3.69$, $p = .001$); right pSTS-I > right FFA in adults ($t(27) = 5.85$, $p < .001$); right pSTS-I > right FFA in children ($t(28) = 2.87$, $p = .004$, one-tailed). Because adults – but not children – showed above-zero selectivity in the bilateral STS-F ROIs, we ran two paired t-tests for adults only. Right pSTS-I selectivity was significantly greater than both the right ($t(27) = 2.61$, $p = .008$, one-tailed) and left STS-F ($t(27) = 3.30$, $p = .002$, one-tailed). Therefore, interaction selectivity was significantly greater in the right pSTS-I than all other ROIs, for both groups.

To compare selectivity in the left pSTS-I with all other regions (except right pSTS-I), in adults (but not children as they did not show above zero selectivity in the left pSTS-I), paired t-tests were performed (Bonferroni corrected $\alpha = .01$). These revealed that interaction selectivity was greater in the left pSTS-I than the right FFA ($t(27) = 3.86$, $p = .001$, one-tailed) and right EBA ($t(27) = 3.05$, $p = .003$, one-tailed), and at an uncorrected threshold in left EBA ($t(27) = 2.01$, $p = .027$, one-tailed) and left STS-F ($t(27) = 1.77$, $p = .044$, one-tailed). However, the trend towards greater selectivity in left pSTS-I than the right STS-F was not significant ($t(27) = 1.24$, $p = .113$, one-tailed). These results demonstrate a moderate trend for greater left pSTS-I selectivity compared to other regions.

*G. Face Selectivity Analyses*

Given the observation for relatively strong interaction selectivity in STS-F, and the close proximity of this region to the pSTS-I – face selective responses were examined. First, face selective responses between right and left pSTS-I ROIs were tested with a 2 x 2 mixed ANOVA (hemisphere x age-group). A marginal main effect of hemisphere ($F$(1,55) = 3.76, $p$ = .057), main effect of age-group ($F$(1,55) = 10.36, $p$ = .002) and no interaction was observed ($F$(1,55) = 0.11, $p$ = .737). These findings show that face selectivity is greater for adults than children in bilateral pSTS-I. A marginal trend is also shown for greater selectivity in the right hemisphere for both groups (although children did not show above zero selectivity in either hemisphere). A similar analysis in the bilateral STS-F revealed no significant ANOVA term, demonstrating that face selectivity is statistically identical between age-groups, in both STS-F ROIs (no main effect of hemisphere: $F$(1,55) = 2.23, $p$ = .141; no main effect of age-group: $F$(1,55) = 0.01, $p$ = .939; no interaction between factors: $F$(1,55) = 0.68, $p$ = .413). Face selectivity trends in bilateral EBA and right FFA were not analysed, but mean selectivity in these regions is plotted below.

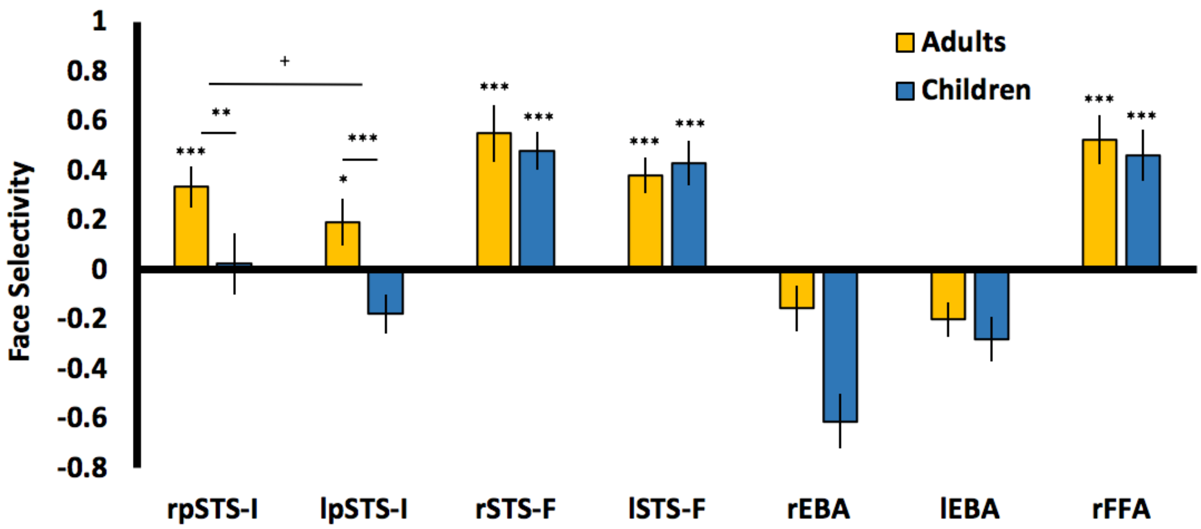


Mean face selectivity values across ROIs. *** = $p$ < .001; ** = $p$ < .01; * = $p$ = .049; $^{+}$ = $p$ = .054. Error bars are SEM. Black circles denote regions that are significantly less face selective than right STS-F for both age-groups.

Next, to determine if face selectivity significantly differed between pSTS-I and STS-F regions, two further ANOVAs were performed. Face selectivity for the right STS-F was significantly greater than the right pSTS-I ($F$(1,55) = 13.64, $p$ = .001), along with a marginal main effect of age-group ($F$(1,55) = 2.88, $p$ = .095), and no interaction between factors ($F$(1,55) = 1.78, $p$ = .187). As with these right hemisphere ROIs, a significant main effect of region ($F$(1,55) = 21.81, $p$ < .001), a marginal main effect of age-group ($F$(1,55) = 3.58, $p$ = .064) was shown between left STS-F and left pSTS-I, however, a significant interaction ($F$(1,55) = 6.20, $p$ = .016) also emerged. Follow up t-tests demonstrated greater face selectivity for faces in the left STS-F than pSTS-I for children ($t$(28) = 4.98, $p$ = .001) – an effect likely driven by 'negative selectivity' for faces in the pSTS-I in children – but this difference was only marginal in adults ($t$(27) = 1.57, $p$ = .064). Therefore, face selectivity in right STS-F was significantly greater than in right pSTS-I, in both age-groups, while this difference was only significant for children in the left hemisphere.

How do these patterns of results relate to interaction selectivity? One general trend is observed: Adults show strongest selectivity in corresponding selective regions (e.g. interactions in pSTS-I and faces in STS-F), but also show weaker responses in the 'other' non-selective STS region (along with other regions). By contrast, children show selective responses in selective regions but not non-selective regions of the STS. Together, these findings suggest that adults recruit wider brain areas during interaction and face perception, whereas children show a greater reliance on selective cortex.

| | | M | SD | t | df | p |
|---|---|---|---|---|---|---|
| rpSTS-I | Adults | 0.37 | 0.44 | 4.01 | 27 | <.001 |
| | Children | 0.02 | 0.67 | 0.20 | 28 | .846 |
| lpSTS-I | Adults | 0.19 | 0.51 | 2.02 | 27 | .053 |
| | Children | -0.18 | 0.43 | -2.23 | 28 | n.s |
| rSTS-F | Adults | 0.55 | 0.61 | 4.78 | 27 | <.001 |
| | Children | 0.48 | 0.41 | 6.36 | 28 | <.001 |
| lSTS-F | Adults | 0.38 | 0.38 | 5.28 | 27 | <.001 |
| | Children | 0.43 | 0.48 | 4.83 | 28 | <.001 |

Descriptive statistics for face selectivity values for adults and children.

*H. Body Selectivity Analyses*

To test whether body selective responses were above-zero in the pSTS-I ROIs, one-sample t-tests (one tailed) were performed. Both adults ($M$ = 0.12, $SD$ = 0.32; $t(27)$ = 1.98, $p$ = .029) and children ($M$ = 0.11, $SD$ = 0.32; $t(28)$ = 1.95, $p$ = .031) showed weak selectivity in the left pSTS-I. By contrast, adults ($M$ = 0.11, $SD$ = 0.32; $t(27)$ = 1.79, $p$ =.043) but not children ($M$ = 0.00, $SD$ = 0.44; $t(28)$ = -0.06, $p$ =.981) showed weak body selectivity in the right pSTS-I.

To compare whether this weak selectivity differed between pSTS-I ROIs, between groups, a 2 x 2 mixed ANOVA (hemisphere x age-group) ANOVA was performed. Neither a main effect of hemisphere ($F(1,55)$ = 1.48, $p$ = .228), main effect of age-group ($F(1,55)$ = 0.60, $p$ = .444), nor significant interaction ($F(1,55)$ = 1.05, $p$ = .309) was found. Therefore, no group or hemisphere differences were observed for weak trends towards body selective responses in the pSTS-I ROIs. Additionally, PSC responses show that both adults and children show strong body responses in bilateral EBA and right FFA (see section E).

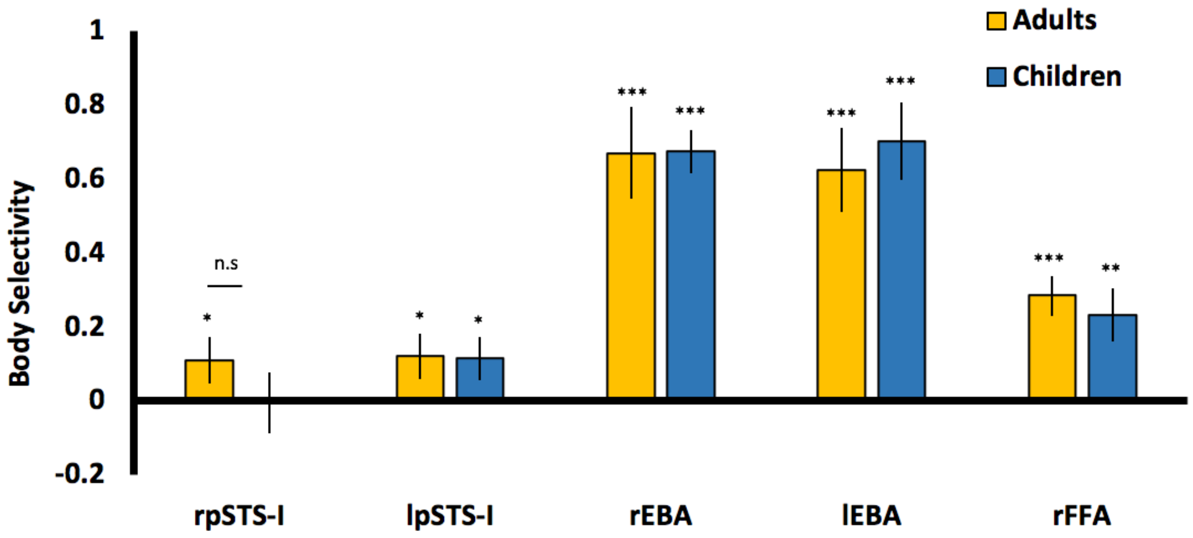


Mean body selectivity values across ROIs. *** = $p$ < .001; ** = $p$ < .01; * = $p$ = .049; + = $p$ = .054. Error bars are SEM.

For comparison to interaction selectivity in pSTS-I, body selectivity in EBA was also analysed with a 2 x 2 mixed ANOVA (hemisphere x age-group). This analysis revealed no significant term (main effect of hemisphere: $F(1,55)$ = 0.22, $p$ = .883; main effect of group: $F(1,55)$ = 0.97, $p$ = .757; interaction term: $F(1,55)$ = 0.38, $p$ = .539), indicating that body selectivity did not differ across hemisphere or age-group. Body selectivity trends in right FFA were not analysed, but mean selectivity in this region is plotted above.

*I. pSTS-I Interaction Selectivity as a Function of ROI Size: Adults vs. Children*

To formally test the apparent age-group differences in interaction selectivity changes with pSTS-I ROI size (i.e. that adults show a linear decrease in selectivity with increasing ROI size, whereas this does not appear to be the case for children), linear regression slopes were calculated for each subject (i.e. the beta coefficient that describes the change in selectivity across ROI size), per ROI and entered into a series of (uncorrected) group tests. One-sample t-tests revealed that adult regression slopes were significantly lower than zero in both the right ($M$ = -0.0015; $SD$ = 0.0032; $t(28)$ = -2.58, $p$ = .007) and left hemisphere ($M$ = -0.0015; $SD$ = 0.0026; $t(28)$ = -3.09, $p$ = .002), but this was not true for children (right pSTS-I: $M$ = -0.0003; $SD$ = 0.0015; $t(30)$ = -1.30, $p$ = .103; left pSTS-I: $M$ = -0.0003; $SD$ = 0.0015; $t(30)$ = -0.91, $p$ = .373). A 2 x 2 mixed ANOVA (hemisphere x age-group) was performed to directly confirm if the two groups differed, and whether this difference was consistent across hemispheres. A main effect of age-group was observed ($F(1,58)$ = 7.05, $p$ = .010), but neither the main effect of hemisphere ($F(1,58)$ = 0.01, $p$ = .912), nor the interaction term was significant ($F(1,58)$ = 0.00, $p$ = .992), indicating that the apparent visual trend was statistically supported in both hemispheres.

*J. Interaction Selectivity Sub-group Analyses*

A one-way ANOVA (with sub-group as the between-group factor) was performed on interaction selectivity values in both the right and left pSTS-I separately, (followed by independent t-tests for marginal and significant ANOVAs, one-tailed, uncorrected $p$ < .05). For the rpSTS-I, a marginal trend was observed ($F(2,39)$ = 2.53, $p$ = .093); adults were significantly more selective for interactions than younger ($t(26)$ = 1.89, $p$ = .035) but not older children ($t(26)$ = 0.49, $p$ = .961), while older children were more selective than younger children ($t(26)$ = 1.78, $p$ = .044). These findings suggest that the marginal trend observed in the main interaction selectivity analysis (i.e. adults > children) is likely driven by weaker selectivity of younger children. Group differences in the lpSTS-I ($F(2,39)$ = 5.90, $p$ = .006), were underscored by greater selectivity for adults compared to both younger ($t(26)$ = 3.41, $p$ = .001) and older children ($t(26)$ = 1.88, $p$ = .035) while a marginal trend for greater selectivity for older than young children was observed ($t(26)$ = 1.59, $p$ = .063).

| | | M | SD |
|---|---|---|---|
| **rpSTS-I** | Adults | 0.40 | 0.31 |
| | 9 – 11 years | 0.39 | 0.35 |
| | 6 – 8 years | 0.11 | 0.47 |
| **lpSTS-I** | Adults | 0.25 | 0.27 |
| | 9 – 11 years | 0.03 | 0.32 |
| | 6 – 8 years | -0.17 | 0.36 |

Descriptive statistics for sub-group interaction selectivity values in the pSTS-I.

*K. pSTS-I Interaction Selectivity as a Function of ROI Size: Sub-group Analyses*

To determine whether similar relationships between pSTS-I ROI size and interaction selectivity as shown for adults and children in the main analysis were also observed across the 3 sub-groups, two analyses were performed on subject's linear regression slopes (i.e. calculated from selectivity values across ROI sizes). One-sample t-tests (one-tailed, uncorrected) revealed that regression slopes were significantly lower than zero in both the right ($M$ = -0.0010; $SD$ = 0.0017; $t$(13) = -2.13, $p$ = .027) and left hemisphere ($M$ = -0.0013; $SD$ = 0.0015; $t$(13) = -3.26, $p$ = .003), but not for either 9 – 11 year olds (right pSTS-I: $M$ = -0.0005; $SD$ = 0.0017; $t$(13) = -1.15, $p$ = .136; left pSTS-I: $M$ = -0.0005; $SD$ = 0.0016; $t$(13) = -1.13, $p$ = .140), or 6 – 8 year olds (right pSTS-I: $M$ = -0.0003; $SD$ = 0.0015; $t$(13) = -0.79, $p$ = .221; left pSTS-I: $M$ = 0.0002; $SD$ = 0.0012; $t$(13) = -0.52, $p$ = .306).

To directly confirm whether these differences were significant between sub-groups, a 2 x 3 mixed ANOVA (hemisphere x sub-group). This revealed a marginal main effect of sub-group ($F$(2,39) = 2.69, $p$ = .080), but neither the main effect of hemisphere ($F$(1,39) = 0.060 $p$ = .807), nor the interaction term was significant ($F$(2,39) = 0.596, $p$ = .556). To follow up the marginal sub-group difference, t-tests (one-tailed, uncorrected) were performed with regression slope values averaged across right and left pSTS-I (as no main effect of hemisphere was observed). Adults were shown to have significantly more negative slopes than 6 – 8 year olds ($t$(26) = -2.00, $p$ = .023), but not 9 – 11 year olds ($t$(26) = -1.15, $p$ = .130), while the two child age-groups did not differ ($t$(26) = 0.70, $p$ = .246).

In summary, results from the one-sample t-tests show that, unlike adults, neither 9 – 11 year olds, nor 6 – 8 year olds show a linear decrease in selectivity with increasing ROI size. Direct comparisons between groups show a statistical difference between adults and 6 – 8 year olds, while the difference between adults and 9 – 11 year olds was not significant, suggesting the possibility of an intermediate level of focal selectivity tuning that is not present in 6 – 8 year olds.

*L. Whole-brain Analyses: Sub-groups*

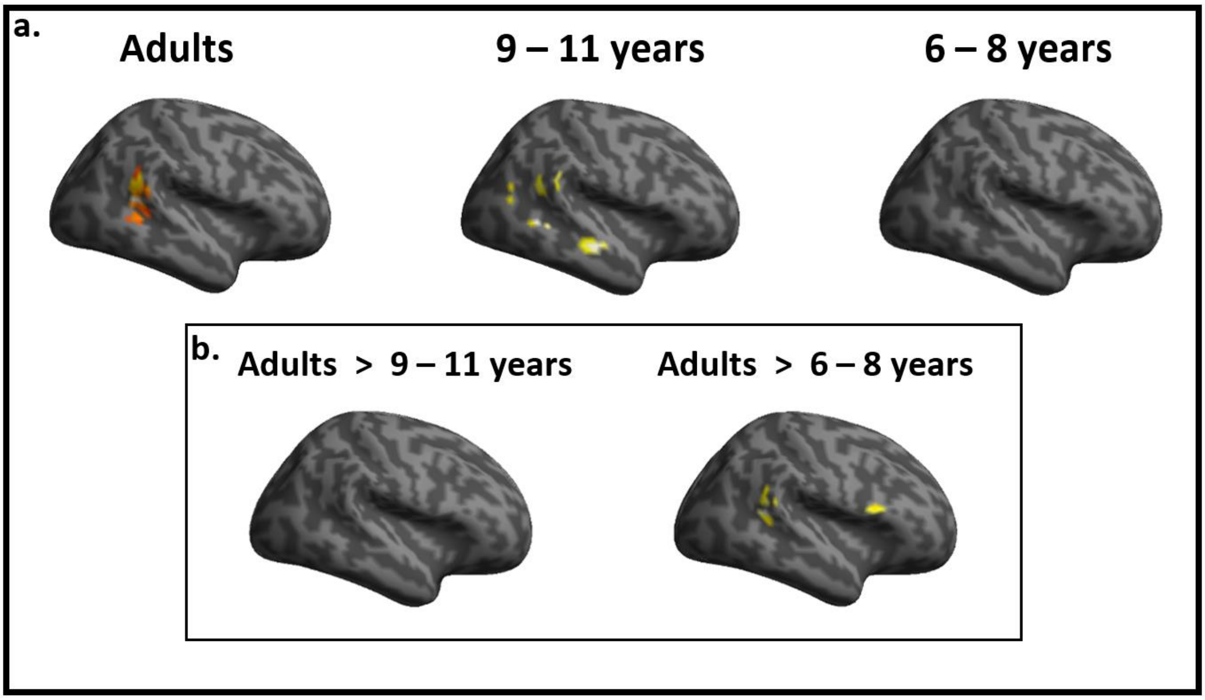


Whole-brain activation for the interaction > non-interaction contrast for the three sub-groups (a), and between-group comparisons (b).

Whole-brain analyses were also conducted for each of the 3 sub-groups, and for between-group comparisons (height-threshold $p$ = .001, FDR cluster-correction $p$ = .05). For the adult sub-group, a similar pattern of activation was shown as in the main analysis with the full set of adult subjects; peak activation in right pSTS and bilateral precuneus (although unlike in the main analysis, no aSTS activation was shown). 9 – 11 years olds showed a very similar pattern of activation to the full group of children in the main analysis with pSTS/pMTG and aSTS activation only, while no activation survived for 6 – 8 year olds. For between-group comparisons, pSTS and IFG activation was shown for the adults > 6 – 8 year olds contrast, but only at a marginal FDR-cluster-correction threshold of $p$ = .09; no activation survived for the adults > 9 – 11 year olds contrast. Overall, these results support the age-related selectivity trends shown for the right pSTS-I.

*M. STS-F Face Selectivity as a Function of ROI Size*

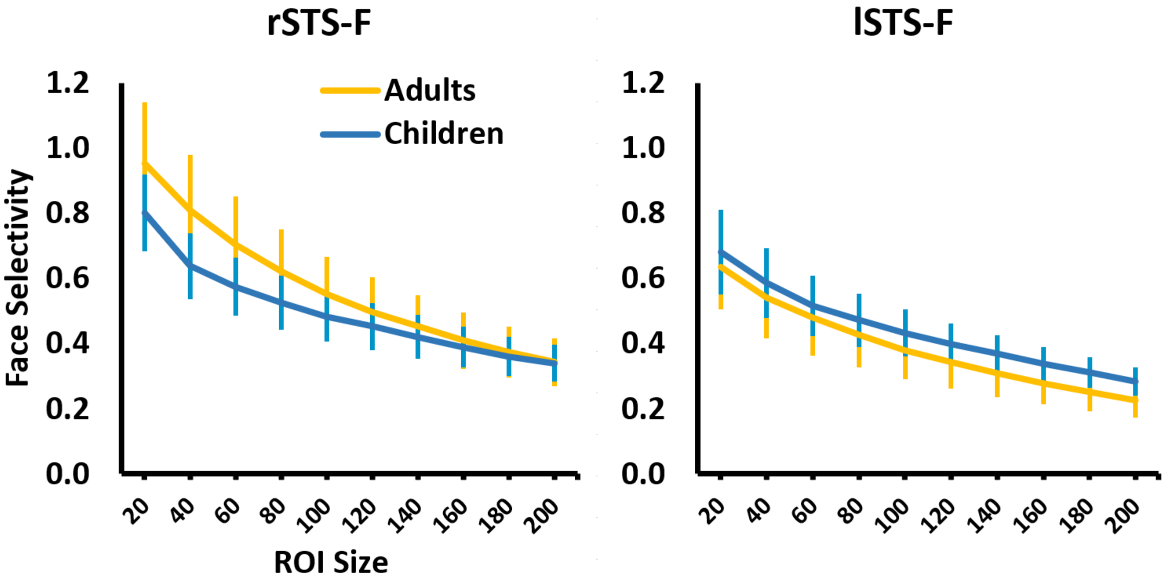


Mean face selectivity plotted as a function of ROI size for adults and children, for the right and left STS-F. Error bars are SEM.

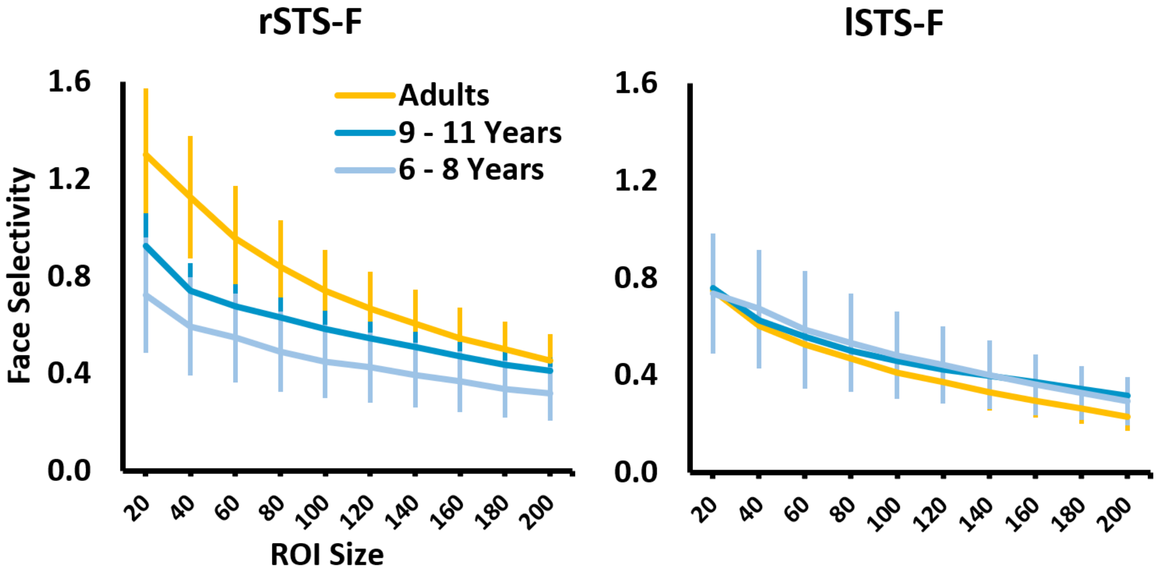


Mean face selectivity plotted as a function of ROI size for the three sub-groups, for the right and left STS-F. Error bars are SEM.

A 2 x 2 ANOVA (hemisphere x age-group) was performed to determine if similar regression slope differences for interaction selectivity in the pSTS-I were shown for face selectivity in STS-F (i.e. all adults vs. all children differences for linear reduction in selectivity with increasing ROI size); no ANOVA term was significant (no main effect of hemisphere: $F(1,55) = 1.70$, $p = .197$; no main effect of age-group: $F(1,55) = 0.75$, $p = .390$); no interaction effect: $F(1,55) = 0.82$, $p = .368$). A similar analysis with the three sub-groups (2 x 3 ANOVA) also revealed no significant term (no main effect of hemisphere: $F(2,37) = 0.97$, $p = .332$; no main effect of age-group: $F(2,37) = 1.34$, $p = .276$); no interaction effect: $F(2,37) = 1.25$, $p = .298$). These findings show that, unlike group differences in focal tuning of interaction selectivity in pSTS-I, no such group differences emerged for face selectivity in the STS-F.
